# Rectangle: robust and scalable multiscale deconvolution informed by single-cell RNA sequencing data

**DOI:** 10.64898/2026.07.07.736950

**Authors:** Bernhard Eder, Irene Rigato, Alexander Dietrich, Lorenzo Merotto, Gregor Sturm, Tim Treis, Markus List, Fabian J. Theis, Francesca Finotello

## Abstract

Bulk RNA-seq enables effective profiling of large cohorts and complex experimental designs, but current single-cell-informed deconvolution methods incompletely resolve closely related cell phenotypes, do not scale efficiently to large single-cell datasets, or fail to account for cellular content not represented in the reference. Here, we present Rectangle, an scverse Python framework for single-cell-informed deconvolution of bulk RNA-seq data. Rectangle combines multiscale deconvolution, capturing cellular composition across multiple resolution levels, with explicit modeling of unknown cellular content. In a diverse, cross-method benchmark, Rectangle achieved consistently strong performance across all evaluated metrics, demonstrating high accuracy, high resolution, low spillover, strong scalability and efficiency, and robustness to unknown cellular content. By bridging the resolution of single-cell transcriptomics with the scale and cost-efficiency of bulk RNA-seq, Rectangle enables cell-type and cell-state profiling at scale, supporting population-scale cellular biomarker discovery and tracking of cellular dynamics in settings impractical for comprehensive single-cell sequencing.

## Background

The structure and function of tissues emerge from the composition and spatial organization of their constituent cells. Quantifying the abundance and localization of cell types is therefore fundamental for understanding development, tissue homeostasis, disease, and responses to therapeutic interventions. *In silico* deconvolution is a computational technique that infers tissue cellular composition directly from bulk RNA sequencing (RNA-seq) data, leveraging cell-type-specific transcriptomic signatures^1,2^. The steep decline in costs has made bulk RNA-seq widely accessible, resulting in thousands of datasets spanning diverse organisms, tissues, and disease contexts, with new data continuously generated in both research and clinical settings worldwide. This growing body of bulk transcriptomic data presents a major opportunity for *in silico* deconvolution to extract cell-type-resolved information at scale.

This potential has been further advanced by recently emerging second-generation deconvolution methods, which learn cell-type-specific transcriptional signatures directly from single-cell RNA-seq (scRNA-seq) data rather than relying on precomputed expression profiles³. Because the single-cell reference can be flexibly defined, second-generation deconvolution is, in principle, applicable to virtually any cell type, tissue, organism, and disease context. However, to fully realize this potential, an ideal deconvolution method should satisfy several key requirements. First, it must enable accurate, fine-grained characterization of tissue composition by reliably distinguishing closely related cell types. Second, to ensure comparability of the inferred cell fractions within and across samples, it should be robust to the presence of “unknown cellular content”, namely, cell types or states not represented in the reference signature matrix. Third, it should be computationally efficient and scalable, enabling routine analysis of the rapidly expanding volumes of single-cell data. More broadly, successful adoption in research and clinical settings requires methods that combine these capabilities with low computational overhead and ease of use. Despite substantial methodological progress, these challenges remain incompletely resolved by existing computational deconvolution approaches^3–6^, limiting their applicability across diverse biological contexts.

To integrate these desired features within a single framework, we developed a novel deconvolution method for the RECursive disenTANGLEment (Rectangle) of bulk RNA-seq samples informed by scRNA-seq data. Rectangle implements a hierarchical strategy for multiscale deconvolution, enabling accurate resolution of fine-grained and closely related cell types. It further leverages a bootstrapping-based pseudobulking strategy to improve robustness to single-cell noise while substantially reducing the computational burden during reference-guided signature building. Additionally, It models unknown cellular content, improving deconvolution robustness and yielding cell-fraction estimates comparable within and across samples. Rectangle is available as an open-source Python package (https://github.com/ComputationalBiomedicineGroup/Rectangle). As part of the scverse ecosystem^7^, Rectangle ensures seamless integration with established data structures and tools, as well as adherence to community standards for documentation, usability, and long-term sustainability.

Using the omnideconv benchmarking framework^6^, we evaluated Rectangle across gold-standard datasets from diverse organisms and tissues, comparing it with eight state-of-the-art methods^8–15^. Across all benchmarks, Rectangle is the only method that consistently combines high accuracy and strong robustness across diverse deconvolution scenarios (lowest cumulative RMSE across all gold-standard RNA-seq mixture datasets) with computational efficiency on a standard laptop (end-to-end analysis based on a ∼150k-cell reference in about one minute). Importantly, Rectangle effectively resolves fine-grained and closely related cell types—a capability it extends to spot-based spatial transcriptomics without requiring specific adaptations. In addition, Rectangle is the only single-cell-informed deconvolution method that estimates and corrects for unknown cellular content, demonstrating its practical relevance in real-world applications such as bulk tumor profiling and longitudinal studies of developing brain organoids. In our comprehensive evaluation, Rectangle improves over the current state of the art, delivering robust and balanced performance across all evaluated categories simultaneously. By jointly addressing key biological and computational challenges, it opens the door to accurate and scalable multiscale deconvolution across diverse biological contexts.

## Results

### Rectangle performs multiscale deconvolution guided by single-cell transcriptomics

Rectangle takes an annotated reference scRNA-seq dataset as input, alongside a bulk RNA-seq dataset to be deconvolved. Its algorithm is articulated into two main steps: *(i)* single-cell-informed creation of the signature matrix and *(ii)* bulk RNA-seq data deconvolution (**Fig. 1**, details in Online Methods).

**Fig. 1.**
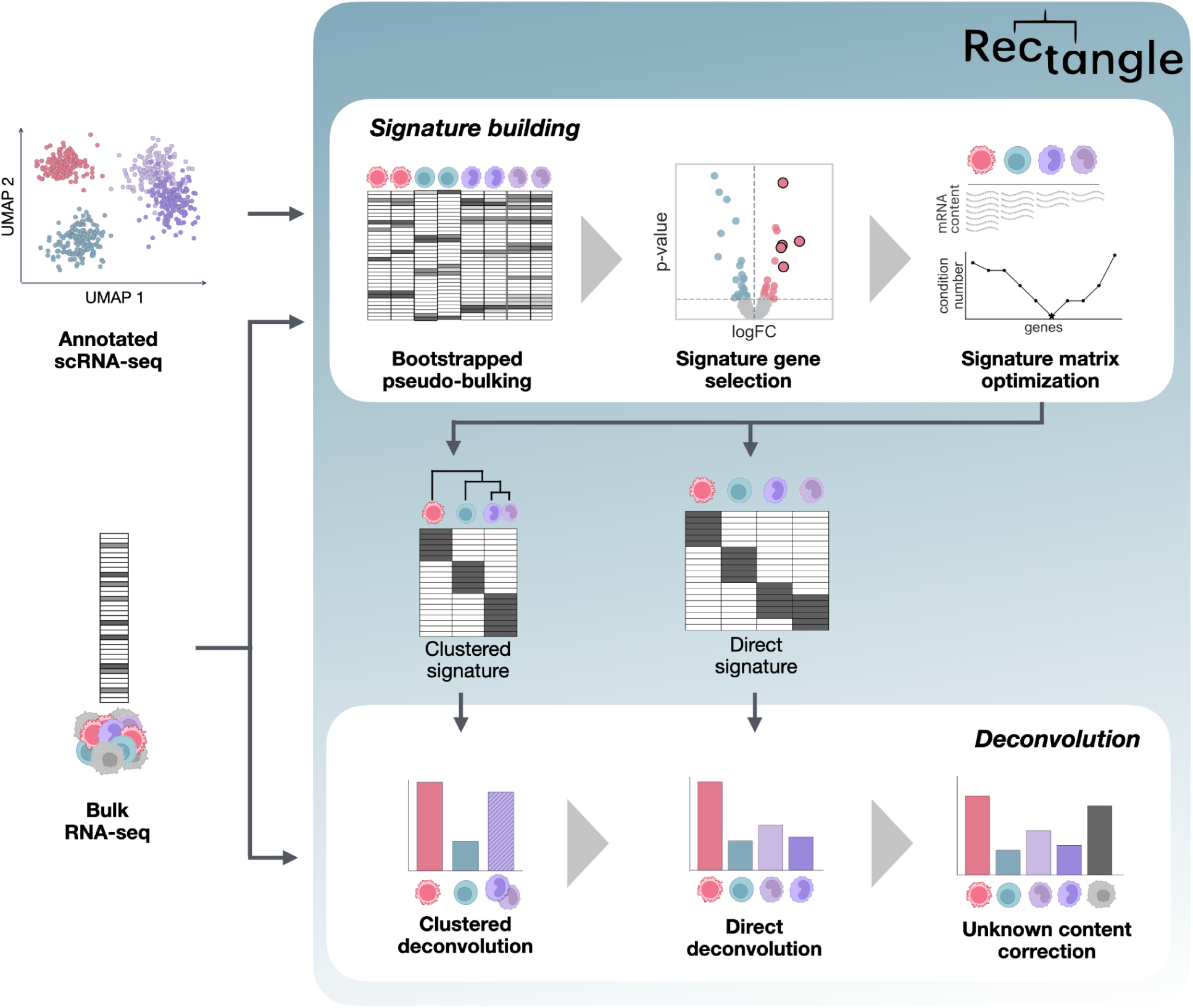
Rectangle performs robust multiscale deconvolution informed by single-cell transcriptomics. Rectangle takes annotated scRNA-seq data and a bulk RNA-seq mixture as input. In the signature-building phase, it performs bootstrapped pseudobulking of the scRNA-seq data, followed by signature-gene selection based on log-fold-change (logFC) and p-value, and signature-matrix optimization (minimizing the condition number and correcting for cell-type-specific mRNA content bias). Based on this, it constructs two signature matrices: a “direct signature”, resolving individual cell types, and a “clustered signature” grouping transcriptionally similar cell types. In the deconvolution phase, Rectangle first uses the clustered signature to deconvolve the bulk data into coarse cell-type estimates. These are then used to constrain a second deconvolution step that leverages the direct signature to resolve individual cell-type fractions. Finally, the resulting estimates are scaled to account for unknown cellular content not represented in the reference.

In the *signature building* phase, Rectangle bootstraps the scRNA-seq data to build multiple pseudobulked, pure expression profiles for every annotated cell type. This bootstrapped pseudobulking process ensures a small memory footprint and runtime, even for very large single-cell references, as well as robustness to data noise and cell-type misannotations. Differential expression (DE) analysis is then performed to identify marker genes, with log-fold-change thresholds adaptively optimized for each cell type using synthetic pseudobulk mixtures. The resulting signature matrix, generated by pseudobulking single-cell profiles, is refined by removing poorly discriminative genes to reduce multicollinearity, and each signature profile is scaled according to cell-type-specific mRNA content to prevent systematic bias in the estimated proportions^1^. This signature building procedure is performed both on the original scRNA-seq annotations (“direct signature”) and on a coarser annotation level obtained by clustering transcriptionally similar cell types (“clustered signature”). The latter aims to identify higher-level marker genes that discriminate high-level lineages and may not be captured via DE analysis of more fine-grained cell subsets.

In the *deconvolution* step, Rectangle quantifies the cellular composition of every bulk RNA-seq sample independently, coupling a quadratic programming (QP)-based deconvolution approach^16^ with the signatures generated in the previous step, while weighting each marker gene with a dampening constant to improve numerical stability^17^. The QP formulation enforces non-negativity of the cell-fraction estimates and constrains their sum to be ≤100%. For highly similar cell types characterized by few or no marker genes, deconvolution is performed in two sequential steps following the hierarchical strategy implemented in Rectangle. First, a QP-based deconvolution on the “clustered signature” provides estimates of coarse cell-group proportions. Second, fine-grained cell-type proportions are inferred using the “direct signature”, guided by the cluster-level estimates. In the QP formulation, cluster proportions are imposed as constraints on the sum of their constituent cell-type fractions within a narrow tolerance interval. This *multiscale* approach restricts the solution space for closely related cell types while enabling fine-resolution estimation. Finally, Rectangle quantifies *unknown cellular content* associated with cell types or states present in the bulk sample but absent from the single-cell reference. This component is estimated from the residual between the observed bulk expression and its reconstruction based on the signature matrix and inferred cell-type proportions. The final cell fractions are rescaled such that they, together with the unknown component, sum to 100%.

### Rectangle uniquely combines accuracy, efficiency, and scalability

We evaluated Rectangle using six validation datasets from mouse and human tissues, for which RNA-seq data and ground-truth cell fractions were available from the same specimens (n=261)^18–23^ (**Suppl. Table 1**). Rectangle showed consistently high overall Pearson’s correlation (0.450–0.941) and low root mean square error (RMSE = 0.056–0.144), with robust performance across cell types, except for myeloid dendritic cells (mDCs, **Suppl. Fig. 1** and **Suppl. Table 2**), which are known to be difficult to quantify due to low abundance and heterogeneous signatures^6,24^. Across all datasets and cell types, Rectangle achieved a mean Pearson’s correlation of 0.804, compared with 0.255-0.754 for the other tested methods. Rectangle was also the most accurate method when considering the cumulative RMSE across all individual cell types and validation datasets (**Fig. 2A**).

**Fig. 2.**
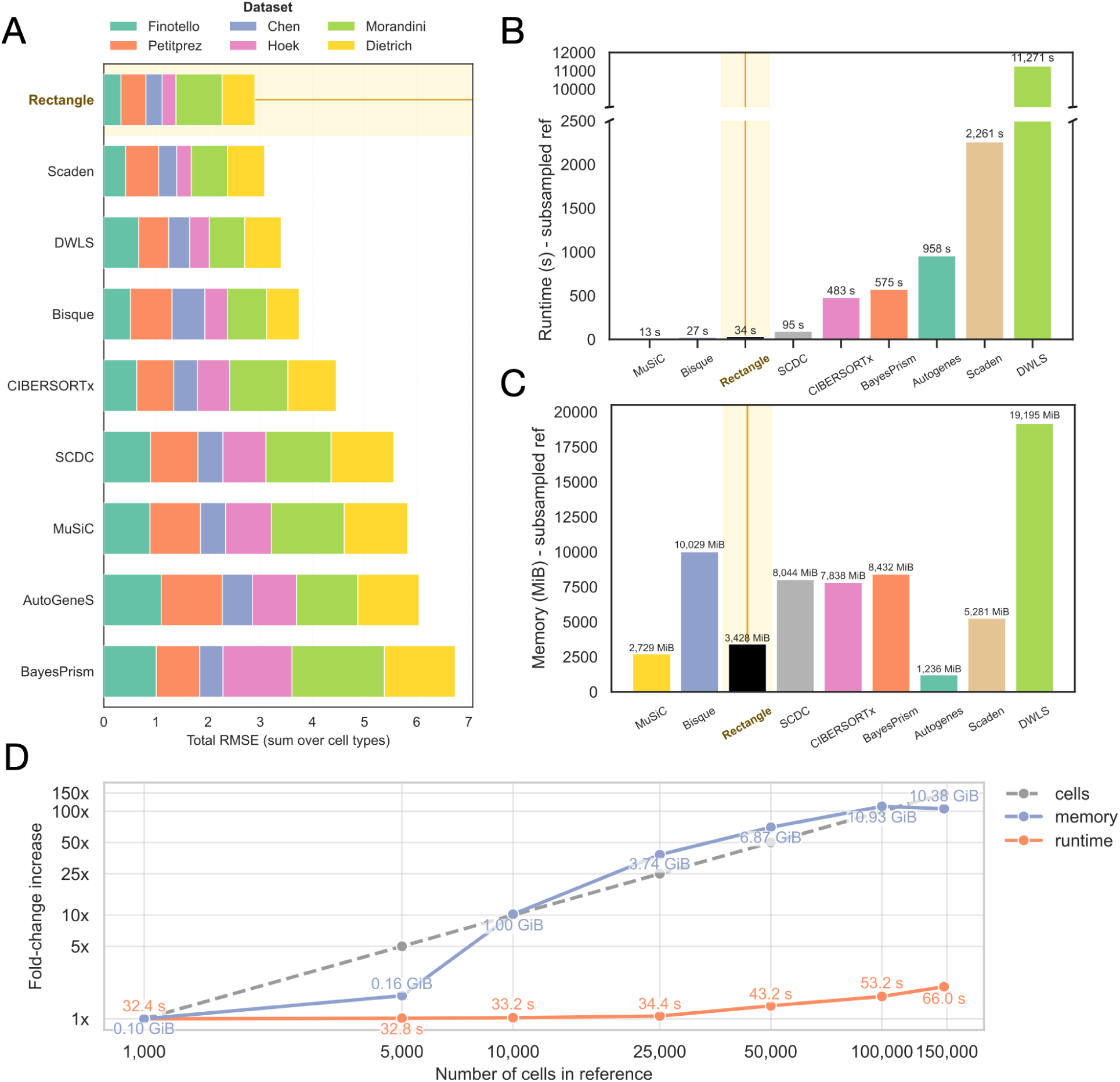
Rectangle combines high accuracy with computational efficiency and scalability. **(A)** Total root-mean-square error (RMSE), summed over all cell types, for each method; stacked bars are colored by dataset. Methods are sorted top-down by increasing total RMSE. Runtime in seconds **(B)** and peak memory usage in MiB **(C)** for each method on the subsampled *Hao* reference. Methods are sorted left-to-right by increasing runtime. **(D)** Scaling of Rectangle deconvolution of the *Finotello* dataset based on reference size: fold-change increase in reference size, runtime, and peak memory relative to the smallest reference, as a function of the number of cells in the reference, and with absolute runtime and memory values indicated.

In light of the continuous growth in scRNA-seq dataset size^25^, the performance of deconvolution tools depends not only on accuracy but also on computational efficiency, which is critical for broad applicability and adoption. We previously showed that accuracy and computational burden often represent opposing trade-offs among state-of-the-art methods^6^. To assess scalability on standard computing hardware, we used an Apple M1 laptop with 8 cores and 32 GB RAM to benchmark the methods for deconvolution of the *Finotello* bulk RNA-seq dataset (n=9)^18^, informed by the *Hao* scRNA-seq dataset (147,391 single cells)^26^, (**Suppl. Table 3**). Only four methods successfully produced estimates, with Rectangle being the fastest (**Suppl. Fig. 2**). Rectangle completed both signature construction and deconvolution in approximately one minute, with runtime dominated by the signature building step (56 s vs. 10 s). We further evaluated scalability using a progressively subsampled version of the *Hao* reference, identifying method-specific limits under our computational settings: Scaden (90k cells), CIBERSORTx (80k), BayesPrism (50k), Bisque (35k), and DWLS (20k). We therefore repeated the benchmark using a uniformly subsampled 20k-cell reference, at which point all methods completed execution (**Fig. 2B,C**). Among these, only Rectangle and MuSiC combined short runtimes (13 s and 34 s, respectively) with low memory requirements (2.7 GiB and 3.4 GiB, respectively).

Via subsampling of the *Hao* dataset, we systematically assessed the behavior of Rectangle across increasing single-cell reference sizes, ranging from 1,000 cells to the full reference (∼150k cells; **Fig. 2D**). Peak memory usage, driven primarily by signature construction, increased approximately linearly with input size, as expected from the underlying data representation and processing steps, rising from 0.10 GiB to 10.38 GiB (104-fold). In contrast, runtime increased only modestly, from 33 s to 92 s (i.e., 3-fold) across this ∼150-fold increase in reference size. This scaling behavior arises from the design of Rectangle: during signature construction, the bootstrapping procedure repeatedly draws a small, constant number of cells and, thus, performance depends only on the number of cell types and genes, and is independent of the size of the single-cell reference. Taken together, these results establish Rectangle as the only method that simultaneously achieves high deconvolution accuracy and low computational burden while maintaining robust scalability across increasing single-cell reference sizes, a capability that is becoming essential as scRNA-seq datasets continue to expand in size.

### Multiscale deconvolution resolves tissue composition at high resolution

Fine-grained deconvolution of tissue complexity depends on methodologies that can resolve transcriptionally similar cell phenotypes, taking full advantage of the resolution provided by annotated single-cell references. To evaluate this capability, “spillover analysis” uses pure bulk samples consisting of a single cell type and assesses whether methods assign the full inferred fraction to the correct cell type^6,24^. Leveraging the *omnideconv* benchmarking framework, we systematically assessed Rectangle’s spillover and compared it with results previously obtained for other state-of-the-art methods^6^. Rectangle exhibited a low spillover effect (average 15.81%), comparable to the best-performing methods, Scaden (13.92%) and CIBERSORTx (15.48%), indicating its ability to minimize erroneous assignment of cell fractions to absent cell types (**Fig. 3A** and **Suppl. Table 4**).

**Fig. 3.**
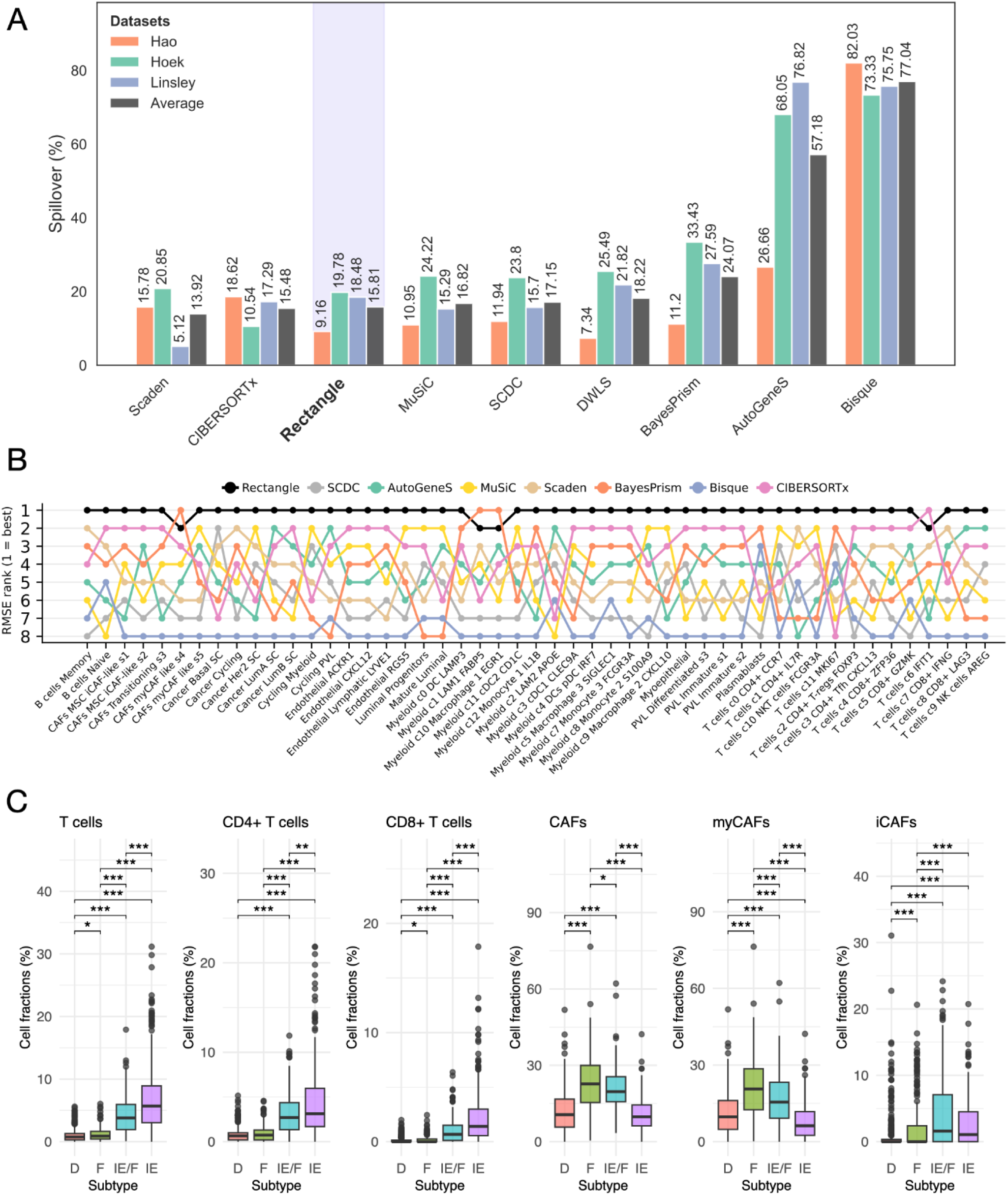
Rectangle enables fine-grained cell-type deconvolution. **(A)** Spillover error (%) quantified for different deconvolution methods across three datasets, together with the average across datasets. Methods are sorted left-to-right by increasing average spillover. **(B)** Bump chart of the method ranks based on root mean square error (RMSE) of each method across 49 fine-grained cell types from the *Wu* dataset. The top-performing method for each cell type appears at the top of the bump chart (rank=1). DWLS is not shown as it failed to complete due to excessive memory consumption. **(C)** Multiscale cell-type fractions deconvolved by Rectangle in breast cancer bulk RNA-seq from The Cancer Genome Atlas, stratified by cancer-immune phenotypes (D, immune-depleted; F, fibrotic; IE/F, immune-enriched fibrotic; IE, immune-enriched), shown for T cells, CD4⁺ T cells, CD8⁺ T cells, cancer-associated fibroblasts (CAFs), myofibroblastic CAFs (myCAFs) and inflammatory CAFs (iCAFs). Significance was assessed by Wilcoxon test (*P < 0.05, **P < 0.01, ***P < 0.001).

We next used the *omnideconv* benchmarking framework to evaluate Rectangle’s performance in a particularly challenging fine-grained deconvolution setting. Using pseudobulk samples generated from breast cancer (*Wu*)^27^ and lung cancer (*Lambrechts*)^28^ scRNA-seq data, which included 49 and 37 annotated cell types, respectively, Rectangle achieved the lowest RMSE for most cell types in both datasets (**Fig. 3B**, **Suppl. Fig. 3**, and **Suppl. Table 5**). These results demonstrate its ability to resolve subtle cellular phenotypes even in highly complex tumor ecosystems. Of note, providing Rectangle with fine-grained labels is also beneficial when the ultimate goal is quantifying coarser cell types, underscoring the importance of its multiscale approach. To demonstrate this, we compared two strategies: *(i)* training Rectangle on fine-grained cell types and aggregating the resulting estimates post hoc into coarse cell-type proportions (i.e., “fine-to-coarse”), and *(ii)* directly training Rectangle using coarse annotations. Tested on the *Wu* and *Lambrechts* pseudobulks, the “fine-to-coarse” strategy consistently yielded more accurate coarse-level estimates, as reflected by a positive reduction in RMSE across all evaluations (4.9-95.3%) (**Suppl. Fig. 4** and **Suppl. Table 6**).

The ability to leverage fine-grained annotations offers a clear advantage for identifying clinically relevant cellular phenotypes that may not align with coarse lineages but instead reflect subtle transcriptional states. We trained Rectangle on the 22 fine-grained cell labels available for the *Wu* dataset to deconvolve 1,091 breast cancer samples from The Cancer Genome Atlas (TCGA)^29^. Notably, for most of these samples, associated cancer-immune phenotype annotations were available: immune-depleted (D, n=400), fibrotic (F, n=215), immune-enriched fibrotic (IE/F, n=180), and immune-enriched (IE, n=273)^30^. These molecular subtypes were originally defined to capture distinct tumor microenvironments (TME) based on immune infiltration and stromal fibrosis, and are associated with differing likelihoods of response to immunotherapy^30^. We examined the deconvolved cell fractions stratified by these four subtypes, considering both fine-grained cell labels and their aggregation into coarser cell types (**Fig. 3C** and **Suppl. Table 7**). Fine- and coarse-grained patterns were concordant in some cases. For example, T-cell subsets exhibited a consistent increase in abundance across the spectrum from immune-depleted to fibrotic, immune-enriched fibrotic, and immune-enriched tumors. In other cases, fine-grained resolution revealed more subtle differences that were not captured at the coarse level, underscoring the added value of fine-grained deconvolution for a clinically-relevant interpretation of the TME. For example, myofibroblastic cancer-associated fibroblasts (myCAFs), which are associated with extracellular matrix remodeling, were enriched in fibrotic tumors (mean in D=11.1%, F=21.2%, IE/F=16.5%, IE=7.9%), whereas inflammatory CAFs (iCAFs), implicated in cytokine signaling and immune modulation, were more abundant in immune-enriched fibrotic tumors (mean in D=1.0%, F=2.2%, IE/F=4.5%, IE=2.8%)^31,32^. Overall, CAFs, instead, presented more similar cell fractions (mean D=12.1%, F=23.4%, IE/F=20.9%, IE=10.7%). Taken together, these results underscore the robustness of Rectangle’s multiscale framework to transcriptionally similar cell types and highlight the value of fine-grained estimates for downstream interpretation. Rectangle enables consistent accuracy propagation across resolutions: improvements at the fine level translate into stable coarse-level estimates, supporting coherent multi-resolution characterization of tissue composition.

### Spatially-resolved deconvolution of fine-grained cell types

While designed for bulk RNA-seq, we reasoned that Rectangle could also be applied to spot-based spatial transcriptomics, where each spot captures a mixed expression profile from multiple cells^2^. To test Rectangle in this setting, we considered a murine brain dataset with 10x Visium spatial transcriptomics data and matched single-nucleus RNA-seq (snRNA-seq) with fine-grained annotations, including region-specific excitatory neuron subtypes. Rectangle could resolve transcriptionally-similar neuronal populations, demonstrating that its multiscale deconvolution accurately mapped these subtypes to their expected anatomical regions (**Fig. 4A**).

**Figure 4.**
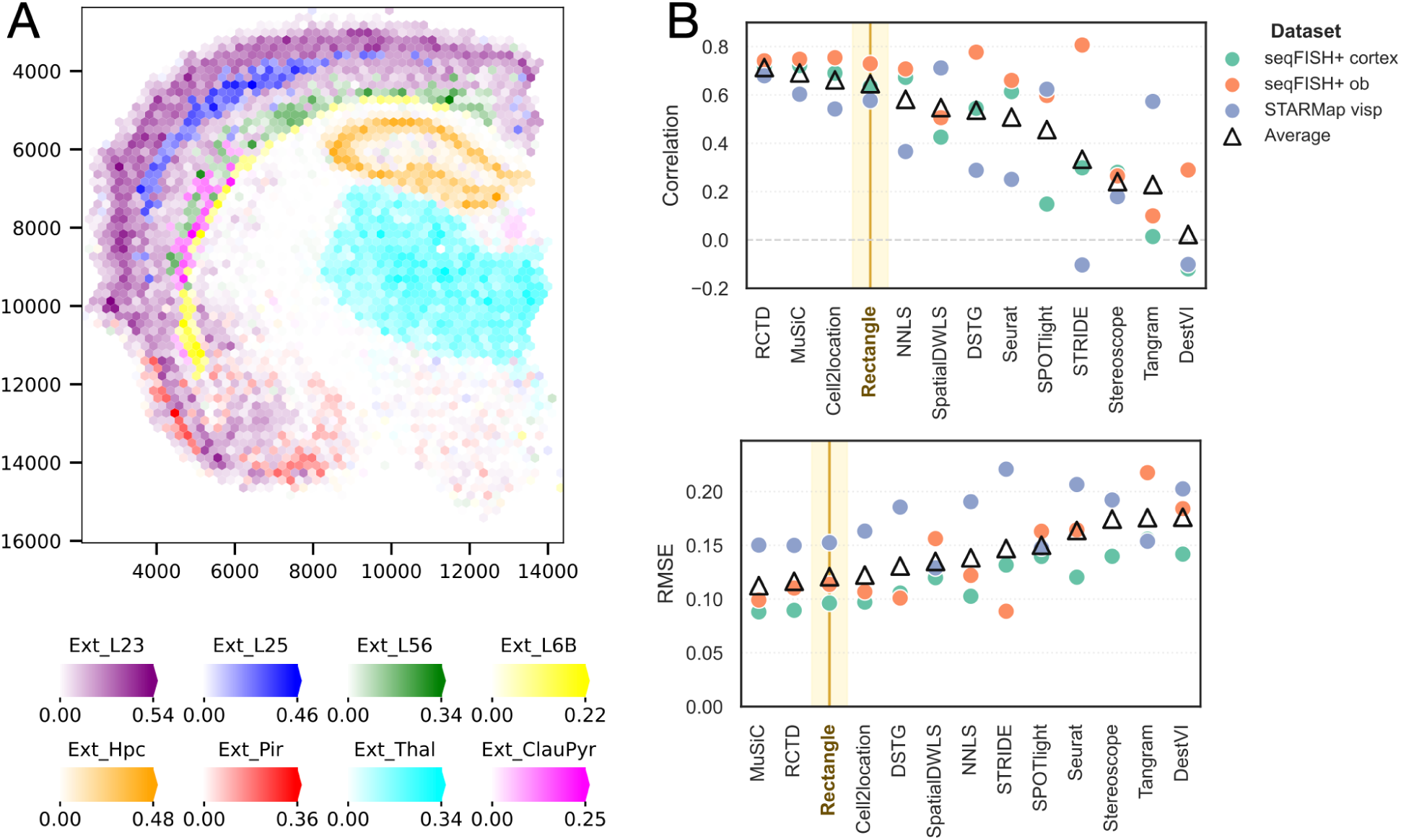
Rectangle extends to spatial deconvolution. **(A)** Spatial map of cell-type proportions predicted by Rectangle across the mouse brain for different excitatory neuron subtypes from the cortical layers (Ext_L23, Ext_L25, Ext_L56, Ext_L6B), hippocampus (Ext_Hpc), piriform cortex (Ext_Pir), thalamus (Ext_Thal), and claustrum (Ext_ClauPyr). The color encodes the inferred cell fraction for the most abundant cell type in every spot. **(B)** Pearson’s correlation (top) and root-mean-square error (RMSE, bottom) between true and estimated cell fractions for different deconvolution methods across three spatial transcriptomics validation datasets. Filled circles denote individual datasets (color), and open triangles denote the average across all three Spotless gold-standard datasets. Methods are ordered by the average value, with correlation sorted descendingly and RMSE ascendingly (top methods on the left).

To quantitatively and systematically assess Rectangle’s performance on spatial deconvolution, we leveraged the established *Spotless* benchmarking framework^33^. Despite not being specifically designed for this task, Rectangle consistently ranked among the best-performing methods based on mean RMSE and Pearson’s correlation across datasets, alongside MuSiC^12^, RCTD^34^, and cell2location^35^ (**Fig. 4B** and **Suppl. Table 8**), which were also identified as top performers in the original benchmarking study^33^. Overall, these results demonstrate that Rectangle extends effectively from bulk RNA-seq to spatial transcriptomics, achieving accurate, fine-grained deconvolution of mixed spatial profiles and competitive performance across established spatial benchmarks.

### Unknown-content estimation resolves hidden malignant and developmental programs

In real deconvolution applications, bulk RNA-seq profiles may contain expression signals from cell phenotypes absent from the scRNA-seq reference and for which no transcriptional signature can be derived, e.g., when cell types are missed for biological or technical reasons^36,37^. This issue can also arise in tumor data, where the marked transcriptional heterogeneity of cancer cells^38^ is difficult to capture in a fixed reference. To provide robust estimates in these settings and express inferred fractions relative to the full cellular composition of the bulk sample, deconvolution methods must estimate and account for such unknown cellular content. Rectangle was specifically designed to address this challenge and is the only single-cell-informed method to offer this feature.

Tested on pseudobulk samples generated with SimBu^39^ and characterized by increasing levels of unknown content, Rectangle was the only method that simultaneously maintained low overall RMSE (0.011-0.047) and high global correlation (0.986-0.998) for reference-covered cell types, even when the unknown component represented a large fraction of the pseudobulk samples (**Fig. 5A,B**). We next evaluated Rectangle on TCGA RNA-seq data from lung squamous cell carcinomas (LUSC) and lung adenocarcinomas (LUAD), using the *Lambrechts* dataset as a reference. Because consensus tumor purity estimates were available for TCGA samples^40^, we could assess whether Rectangle’s unknown-content estimates captured a tumor-derived signal not represented in the reference. When trained only on the non-tumoral compartment of the *Lambrechts* dataset, Rectangle recovered a substantial fraction of this hidden tumor-specific signal, as shown by the agreement between the estimated unknown content and per-sample tumor purity (r=0.612, RMSE=0.285, **Fig. 5C**). When trained on the full *Lambrechts* dataset, the estimated unknown content remained non-negligible, ranging from 0 to 36.4% across samples, with an average of 5.0%. This suggests that Rectangle captured a residual tumor-specific signal reflecting inter-tumor transcriptional heterogeneity not fully represented by the *Lambrechts* reference. Accordingly, combining the tumor-cell with unknown content fractions estimated by Rectangle improved the quantification of tumor purity relative to the tumor estimate alone, both in terms of global Pearson’s correlation (0.772 vs. 0.646) and RMSE (0.112 vs. 0.141) (**Fig. 5C**).

**Fig. 5.**
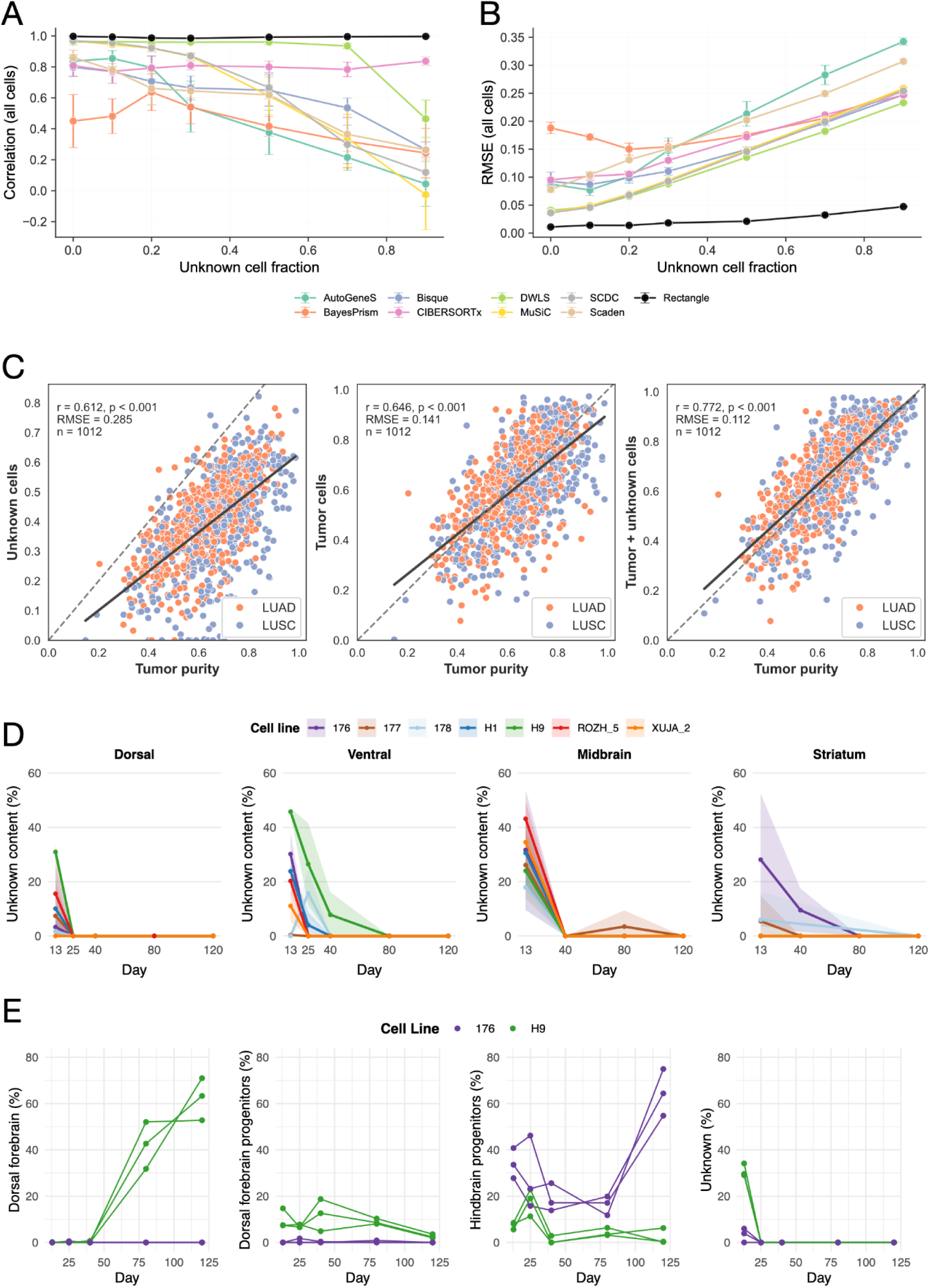
Quantification of unknown cellular content expands deconvolution beyond reference-defined cell types. Global Pearson’s correlation **(A)** and root-mean-square error (RMSE) **(B)** between true and estimated fractions for the reference-covered cell types obtained by different deconvolution methods applied to lung cancer pseudobulks containing increasing unknown tumor-cell content. **(C)** Scatter plots comparing Rectangle’s estimates with tumor purity for lung adenocarcinoma (LUAD, orange) and lung squamous cell carcinoma (LUSC, blue) samples from The Cancer Genome Atlas. Left, unknown content estimated when no scRNA-seq reference for tumor cells is supplied; middle, tumor-cell fraction estimated when a tumor-cell reference is provided; right, sum of the estimated tumor-cell fraction and unknown content when a tumor-cell reference is provided. Dashed lines denote the identity (y=x); solid lines show the linear fit. Pearson correlation (r), RMSE and sample number (n) are indicated. **(D)** Rectangle’s estimated unknown cellular content over time in bulk RNA-seq data from developing brain organoids, across four differentiation protocols (dorsal forebrain, ventral forebrain, midbrain, striatum) and seven cell lines (color). Shaded areas denote the standard deviation across replicates. **(E)** Time-resolved cell fractions deconvolved by Rectangle for the H9 and 176 cell lines during dorsal forebrain differentiation, shown for selected cell types (dorsal forebrain, dorsal forebrain progenitors, hindbrain progenitors, and unknown content).

To further show the relevance of Rectangle’s “unknown content” estimation module, we applied it to bulk RNA-seq data from developing brain organoids. The organoids were generated from multiple pluripotent cell lines using four protocols, each aimed at generating the cellular composition of different brain regions: dorsal and ventral forebrain, midbrain, and striatum^41^. In the original study, the bulk data were generated at different time points (days 13/16, 25, 40, 80, and 120) to track the development of brain tissue. In contrast, annotated scRNA-seq data were available only for day 120, representing the final cellular phenotypes. Thus, in the deconvolution of the time-series bulk data, we would expect some unknown cellular content at the early time points, representing early progenitors or not fully committed cell types, which are not captured by the single-cell reference. Indeed, Rectangle deconvolution showed a larger unknown content in the first time point across all protocols, cell lines, and biological replicates (**Fig. 5D**) (0.0%-53.7%). The only exception is represented by the 178 cell line in the ventral protocol, which showed a peak in correspondence with the second time point (day 25). Because Rectangle estimates unknown content from discrepancies between the reconstructed and observed bulk profiles, genes with high reconstruction error can help pinpoint transcriptional programs present in the bulk data but absent from the reference. Rectangle reports this per-bulk reconstruction error for each gene and sample, thereby facilitating the investigation of the hidden component. By analyzing the top 10 genes with the largest reconstruction error at days 13/16 across all cell lines and protocols, we identified a shared early program characterized by early neurogenic differentiation or neural precursor-state transitions (SOX11, PRTG), cytoskeletal and microtubule remodeling (TUBA1B, TUBB, TUBB2B), proliferation-associated chromatin and cell-cycle regulation (NUCKS1, NAP1L1), increased translational/ribosomal activity (RPS29, RPL31), and proteostasis/stress-response regulation (HSP90AA1). Together, these reconstruction-error signatures indicate that the unknown component quantified by Rectangle quantitatively disentangles a conserved, early progenitor-like signature in developing brain organoids.

By inspecting the full set of cell types deconvolved by Rectangle across the 363 analyzed bulk samples (**Suppl. Table 9**), we can gain insight into brain organoid development and investigate cellular shifts associated with successful or unsuccessful reprogramming. For example, in the original study, the H9 cell line was identified as a prototypical successful line for the dorsal protocol, whereas the 176 cell line failed to recapitulate the expected cell phenotypes. Consistently, the cellular compositions inferred by Rectangle across time points and biological replicates (**Fig. 5E** and **Supplementary Fig. 5**) revealed clear differences between these two lines. At the first time point, the 176 cell line exhibited lower unknown content (0.0-0.6%) than the H9 cell line (29.0-34.1%). By the final time point, H9 showed successful conversion toward dorsal forebrain cells (56.4-73.0%), a pattern already broadly visible at day 80 (39.9-62.4%). In contrast, the 176 cell line did not acquire dorsal-cell phenotypes and was instead dominated by hindbrain-associated cell phenotypes. Together, these results establish unknown-content estimation and dissection as a key feature for deconvolution in real biological settings. This allows Rectangle to move beyond reference-matched cell-type quantification and recover hidden tumor programs, early developmental states, and lineage deviations that would otherwise remain unexplained.

## Discussion

Cell-type deconvolution is becoming central to the interpretation of large transcriptomic datasets, but its broader application requires methods that can jointly resolve closely related cellular states, scale to increasingly large datasets, and remain informative when references are incomplete. Here, we showed that Rectangle addresses these challenges through a multiscale framework that combines accurate and scalable deconvolution with explicit detection of transcriptional content not captured by the reference. In our comprehensive assessment, Rectangle achieved the most consistently robust and balanced performance across all evaluated metrics (**Figure 6**).

**Fig. 6.**
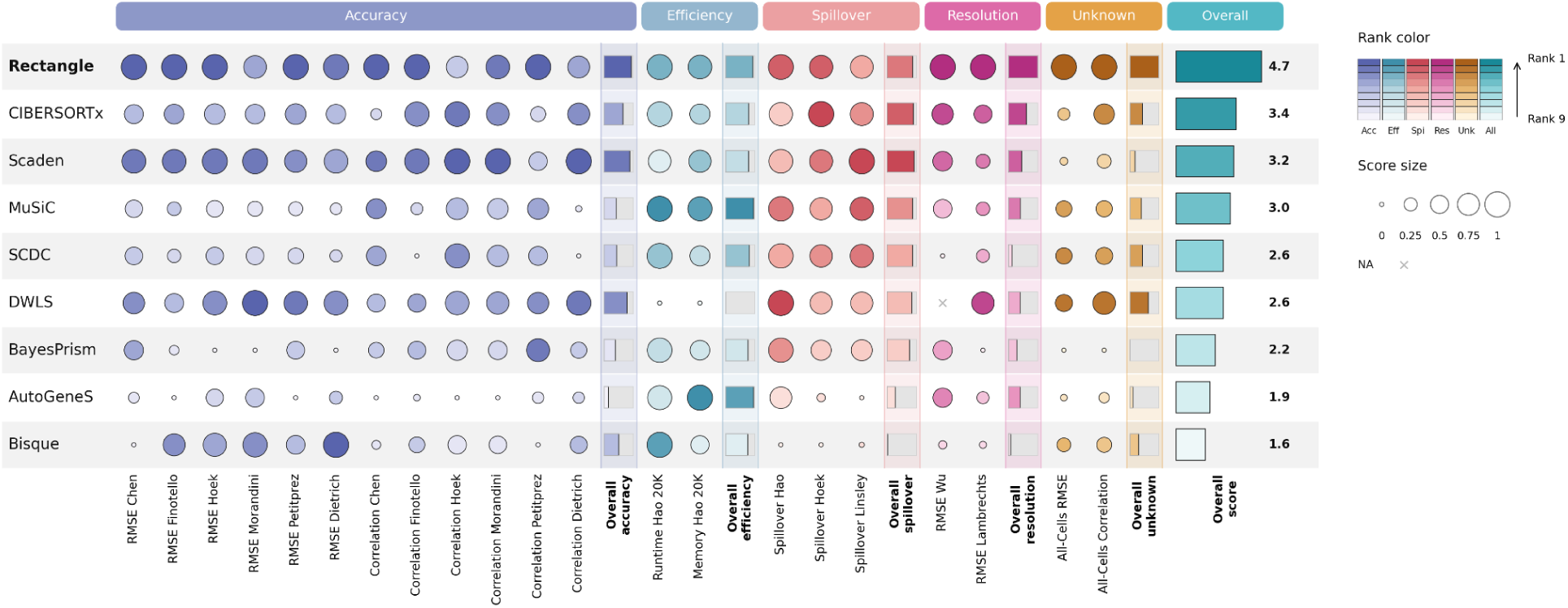
Rectangle’s top overall performance is grounded in robust results across all benchmark categories. Integrated summary combining all benchmarks into five performance categories: deconvolution accuracy, computational efficiency, spillover error, cell-type resolution, and robustness to unknown content. Each original metric was min-max normalized in [0, 1], with 1 denoting the best method for each category (details in Online Methods). Marker size encodes the normalized score (the larger the better), and color encodes the within-category rank, from rank 1 (best) to rank 9 (worst). The “overall” bar per category is the mean of its normalized metrics. The rightmost column gives the overall score, by which methods are ordered. Missing values are marked with “x”.

Rectangle implements a multiscale approach to cell-type and cell-state quantification, which builds on the principle that hierarchy-guided deconvolution can mitigate multicollinearity among transcriptionally similar phenotypes^12^. This strategy is particularly important in modern single-cell references, where annotations increasingly distinguish closely related cell states that may differ by subtle transcriptional programs. Across benchmark datasets, Rectangle maintained high accuracy while limiting spillover between transcriptionally similar populations, demonstrating its ability to resolve extremely fine-grained cell states in both complex tumor microenvironments and the anatomically structured mouse brain. Importantly, this fine-grained resolution did not compromise coarser-level interpretation. In fact, estimates obtained from fine-grained labels could be aggregated into accurate broad cell-type fractions, outperforming direct coarse deconvolution—a behavior also observed for other deconvolution methods^6,9^.

Rectangle also addresses a practical bottleneck for the field: scalability. By combining pseudobulk-based signature construction, sparse data structures, and extensive parallelization, Rectangle achieves low runtime and memory usage, enabling scalable deconvolution on a standard laptop without compromising accuracy. This balance is becoming essential as single-cell references continue to increase in size and complexity^25,42,43^. Our study and previous benchmarks^4,6^ showed that existing approaches typically trade off deconvolution accuracy against computational efficiency. Instead, Rectangle uniquely combines both properties, offering a practical framework for atlas-scale transcriptomic deconvolution that preserves fine-grained accuracy while remaining computationally efficient. Beyond performance, Rectangle was designed for broad usability, with straightforward installation, extensive documentation and interoperability across major single-cell and spatial transcriptomics ecosystems. In Python, Rectangle integrates with the scverse ecosystem through native support for AnnData^44^ and interoperability with SpatialData^45^. In R, it is accessible through omnideconv^6^ and spacedeconv^46^.

Although Rectangle was developed for bulk RNA-seq, our results indicate that its algorithmic strategy extends naturally to spot-based spatial transcriptomics. Without any adaptations to spatial-specific features, Rectangle recovered anatomically coherent fine-grained brain populations and ranked among the top-performing methods in our spatial deconvolution assessment supported by the Spotless framework^33^. This suggests that Rectangle can perform spatial deconvolution in contexts where tissue architecture carries biological information, enabling the characterization of spatially organized cellular niches and regional tissue domains. Here, we focused on 10x Visium spot-based data due to its rapid adoption and the large volume of available datasets^47^; however, our framework readily extends to high-resolution spatial technologies in which near-cellular measurements are aggregated into larger, deconvolvable spatial bins^48^. Future work could incorporate analytical modules tailored to spatial data, such as enrichment-based restriction of candidate cell types per spot/tile^49^ and spot-level modeling of cell abundance and tissue density^35,46^.

A key contribution of Rectangle is the explicit estimation of unknown transcriptional content. In real-world applications, reference incompleteness is common: relevant cell types or states may be absent from the single-cell reference^36,37^ because of experimental design, technical sampling biases, or context-dependent transcriptional shifts. Tumors represent an especially important case, given the extreme heterogeneity of malignant cells^38^, the limited availability of exactly matched single-cell references, and the clinical relevance of cellular biomarkers defined within the TME, as exemplified by the prognostic value of the tumor immune contexture^50^. For transcriptomic deconvolution, this creates an additional requirement: estimated fractions should reflect the proportion of each cell type in the entire sample, not only among reference-defined populations. If tumor cells or other major components are missing from the reference, known cell types may be artificially rescaled to fill the unexplained fraction, leading to misleading cellular biomarkers in cohort-scale analyses. This need was recognized by first-generation methods such as quanTIseq^18^ and EPIC^51^, which included cancer-cell or unassigned fractions, but has been largely overlooked by second-generation, single-cell-informed methods that typically assume complete reference coverage. Rectangle addresses this issue by explicitly quantifying the fraction of the bulk transcriptome that cannot be explained by the provided reference. In TCGA lung cancer samples, this unknown component captured a tumor-associated signal not represented in the non-tumoral reference and improved tumor purity estimation when combined with reference-guided tumor-cell estimates. This feature was also relevant in the analysis of developing brain organoids. By deconvolving time-course bulk data against a single-cell reference built exclusively from mature cells, Rectangle identified a fraction of early samples as states not represented in the reference, consistent with immature progenitor populations, with the associated genes defining a coherent early developmental program. This analysis further revealed cellular dynamics associated with successful and unsuccessful brain organoid differentiation trajectories, consistent with the molecular findings reported in the original study^41^. Together, these examples show that Rectangle can transform unexplained residual expression into an interpretable signal, revealing “hidden” cell types and their associated transcriptional programs.

More broadly, the brain organoid analysis illustrates a powerful methodological framework that is readily extendable to other contexts: transcriptomic-based cellular profiling at scale. A focused single-cell reference can be integrated with large bulk RNA-seq collections to infer cellular composition across samples, conditions, and time points, combining the resolution of single-cell transcriptomics with the scalability, lower cost, and unbiased nature of bulk RNA-seq. This enables the large sample sizes required for clinically relevant biomarker discovery and transforms bulk transcriptomic datasets into digitally resolved cellular compositions.

Rectangle is robust to noise and annotation errors in scRNA-seq references through bootstrapping and multiscale, hierarchical deconvolution. Nevertheless, some limitations remain due to its data-driven algorithmic approach, which inevitably depends on the quality and completeness of the single-cell reference. Although we did not perform extensive reference curation in this study to avoid dataset-specific engineering, we recommend careful inspection of the reference in real applications, particularly for highly similar cell types whose boundaries may be difficult to define even at single-cell resolution. As a practical guideline, clusters annotated primarily by cell-cycle activity should generally be removed, as they can obscure the identification of truly cell-type-specific markers shared with their less proliferative counterparts. Similarly, cell types having too few single-cell profiles in the references should be excluded from deconvolution analysis. Future developments will include preprocessing modules to automatically flag hard-to-resolve phenotypes to guide these reference-refinement steps, alongside confidence intervals and statistical significance estimates for the inferred cell fractions, enabling more rigorous uncertainty quantification for each deconvolution result.

In conclusion, by integrating the cellular resolution of single-cell transcriptomics with the scale and cost-efficiency of bulk RNA-seq, Rectangle provides a foundation for multiscale analysis of cellular composition across large cohorts, time series, and high-throughput screening and perturbation studies. This framework establishes a path toward population-scale cellular profiling for identifying clinically relevant biomarkers, stratifying patients, and tracking cell-type and cell-state dynamics in settings that are impractical for comprehensive single-cell sequencing.

## Data availability

The full set of datasets used in this study is reported in **Suppl. Table 1**. No other data were generated within this study.

## Code availability

Rectangle code is available at: https://github.com/ComputationalBiomedicineGroup/Rectangle. Rectangle documentation can be accessed from: https://rectanglepy.readthedocs.io/. All scripts to reproduce the analyses of this study are available from: https://github.com/ComputationalBiomedicineGroup/rectangle_paper.

## Acknowledgements

This work was supported by the Vice-Rectorate for Research and Förderkreis 1669 of the University of Innsbruck. This article is based upon work from COST Action Mye-InfoBank (CA20117) and COST Innovators Grant Omnicellscope (IG20117), supported by COST (European Cooperation in Science and Technology). FF was supported by the Austrian Science Fund (FWF) (Grant-DOI 10.55776/FG25 and 10.55776/PAT5895324). The computational results presented here have been achieved in part using the LEO HPC infrastructure of the University of Innsbruck, and the de.NBI Cloud within the German Network for Bioinformatics Infrastructure (de.NBI). The results shown here are in part based upon data generated by TCGA Research Network (https://cancergenome.nih.gov). The authors are grateful to Dr. Christopher Esk for insightful discussions on the brain organoid analyses.

## Conflicts of interest

FF consults for iOnctura. G.S. is an employee of Boehringer Ingelheim International Pharma GmbH & Co KG, Biberach, Germany. F.J.T. consults for Immunai, CytoReason, Valinor Industries, Bioturing and Phylo Inc., and has ownership interest in RN.AI Therapeutics, Dermagnostix, and Cellarity.

## Online Methods

### Rectangle algorithm

The Rectangle algorithm is packaged as an open-source Python library. The complete source code, documentation, and installation instructions, are available at https://github.com/ComputationalBiomedicineGroup/Rectangle. Rectangle takes as input a tag-based scRNA-seq reference dataset, provided either as a dense or sparse gene-count matrix plus cell-type labels for each individual cell, and a bulk RNA-seq dataset to be deconvolved, in the form of a transcripts per million (TPM)-normalized gene expression matrix. For deconvolution of spot-based transcriptomics, counts per million (CPM)-normalized data are expected. Rectangle’s algorithm operates in two main phases: *(i)* single-cell-informed creation of the signature matrix and *(ii)* bulk RNA-seq data deconvolution.

In the *signature building* phase, Rectangle bootstraps the scRNA-seq data to build pseudobulk expression profiles for every annotated cell type. For each cell type, it randomly samples 500 cells with replacement from all cells of that type and sums their gene-count profiles to produce a “pure” pseudobulk sample. By default, this bootstrapped pseudobulking process is repeated seven times per cell type, although this parameter can be adjusted by the user. This strategy ensures a small memory footprint and runtime even in the case of very large single-cell references, as well as robustness to data noise and cell-type misannotations.

Differential expression (DE) analysis is then performed with pyDESeq2^1^ to identify cell-type-specific marker genes as a basis for signature building, retaining only genes that are upregulated in the target cell type, i.e., that have a positive log_2_-fold change (LFC). DE analysis is performed as a one-versus-all comparison: the bootstrapped pseudobulk samples of the selected cell type are labeled as the target condition, while the bootstrapped samples of all remaining cell types form the reference condition. To identify DE genes, the false discovery rate (FDR)-adjusted p-value cut-off is fixed at 0.05, while the LFC threshold is adaptively determined via a grid-search approach, informed by synthetic pseudobulk cell mixtures. Specifically, Rectangle searches over a list of positive LFC values and, for each candidate LFC threshold, it derives the resulting DE genes and an interim signature matrix. This signature matrix is used to deconvolve a set of 50 synthetic pseudobulk samples, generated by randomly drawing without replacement 0-50 cells per cell type from the scRNA-seq reference, summing up their expression profiles. The upper limit of 50 cells per cell type is used by default to generate heterogeneous synthetic mixtures while keeping the optimization computationally efficient, but this parameter can be adjusted by the user. Allowing zero sampled cells creates pseudobulks in which individual cell types are absent, which is important for evaluating whether a candidate signature can recover both present and missing populations. The optimal LFC threshold per cell type is selected by maximising the Pearson’s correlation coefficient between the estimated and true cell fractions across all pseudobulk mixtures, with root mean square error (RMSE) used as a tiebreaker.

The identified DE genes are used to assemble a preliminary signature matrix by pseudobulking all single-cell profiles for each cell type. The resulting profiles are normalized to CPM to match the scaling of bulk RNA-seq data (TPM) or spatial transcriptomics data (CPM), ensuring comparable correction for library size and gene-length bias. This signature is further optimized by removing genes with low discriminative power that introduce collinearity, thereby minimizing its condition number, a measure of numerical stability and robustness to noise. Rectangle achieves this by constructing multiple candidate signature matrices, each containing a different number of top-ranked marker genes (sorted by LFC) per cell type (by default, ranging from 20 to 80). For each candidate signature matrix, QR decomposition is first applied to factor the matrix into an orthogonal component Q and an upper-triangular component R. The condition number is then computed on the R factor using the matrix 1-norm, and the matrix with the lowest condition number is selected. Moreover, each signature profile is scaled by a factor reflecting cell-type-specific mRNA content to prevent systematic bias in the estimated proportions^2^. Rectangle estimates this cell-type-specific mRNA bias factor from the scRNA-seq reference, considering the mean number of expressed genes per cell as a proxy for mRNA abundance, as proposed by Dietrich et al.^3^. Specifically, it counts the number of genes with nonzero expression in each cell, averages these counts within each cell type, and normalizes the resulting averages by the smallest cell-type-specific value. Thus, the cell type with the lowest inferred mRNA content receives a bias factor of 1, while all other cell types are scaled relative to it.

The signature building and optimization steps are performed both on the original scRNA-seq annotations (“direct signature”) and on a coarser annotation level obtained by clustering transcriptionally similar cell types (“clustered signature”). The latter aims to identify higher-level marker genes that discriminate related lineages and may not be captured by DE analysis of more fine-grained cell subsets. To construct the clustered signature, Rectangle first applies hierarchical agglomerative clustering to the cell types represented in the direct signature. Specifically, the marker gene expression profiles in the direct signature are log-transformed and clustered using complete linkage with Euclidean distance. The optimal number of clusters is selected by maximizing the silhouette score over a narrow range close to the original number of cell types, allowing at most four merged cell-type groups. This constraint prevents excessive aggregation of transcriptionally distinct cell types while still permitting closely related annotations to be combined. Every cell type that is the sole member of its cluster retains its original annotation, while cell types assigned to a shared cluster are reannotated with a common cluster label. These cluster-level labels replace the original cell-type labels for the corresponding single cells, and the full signature-building procedure is then repeated using this clustered annotation.

In the *deconvolution* phase, Rectangle independently quantifies the cellular composition of each bulk RNA-seq sample by coupling a quadratic programming (QP)-based deconvolution approach^4^ with the signatures generated in the previous step. For each bulk sample, Rectangle estimates cell fractions by minimizing the weighted squared difference between the observed bulk expression and the expression reconstructed from the signature matrix: 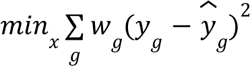 subject to *x _i_* ≥ 0 and 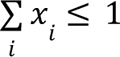. Here, *x _i_* is the estimated fraction of cell type *i*, *y _g_* is the observed expression of gene *g, ŷ_g_* is the reconstructed expression of gene *g*, and *w _g_* is the dampened marker-gene weight^5^. The latter assigns each gene a contribution weight following the dampened weighted least squares (DWLS) principle^5^. Instead of minimizing only absolute reconstruction error, genes are weighted inversely to their squared reconstructed expression, giving lowly expressed but informative marker genes greater influence. The QP formulation enforces non-negativity of cell-fraction estimates and constrains their sum to be ≤100%.

For highly similar cell types characterized by ≤ 20 marker genes, deconvolution is performed in two sequential steps following the hierarchical strategy implemented in Rectangle. First, a QP-based deconvolution on the “clustered signature” provides estimates of broad cell-group proportions. Second, fine-grained cell-type proportions are inferred using the “direct signature”, guided by the cluster-level estimates. In the QP formulation, cluster proportions are imposed as constraints on the sum of their constituent cell-type fractions. For each cluster (k), the estimated fine-grained fractions are constrained according to 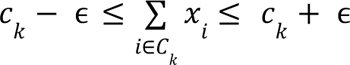 where *c _k_* is the cluster-level proportion estimated from the clustered signature, *c _k_* is the set of cell types belonging to that cluster, *x _i_* is the fraction of cell type *i*, and a tolerance of ɛ = 0.03. This value was set to allow moderate flexibility around the cluster-level estimate while still preventing the second deconvolution step from substantially redistributing mass across broad cell groups. This restricts the solution space for closely related cell types while enabling fine-resolution estimation. Finally, unknown cellular content is estimated as the fraction of total bulk expression not explained by the reconstructed profile. The reconstructed expression of gene *g* is defined as 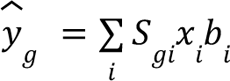 where *S _gi_* is the signature expression of gene *g* in cell type *i*, *x i* is the inferred fraction of cell type *i*, and *b i* is the corresponding mRNA bias factor. Unknown cellular content is then estimated as 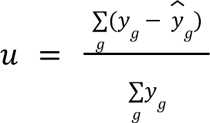, with negative values set to zero. The final known cell-type fractions are then rescaled to sum to 1 − *u*, and the unknown component is reported as *u*.

### Bulk RNA-seq deconvolution benchmarking

All the single-cell references and bulk RNA-seq datasets with ground-truth cell fractions from the omnideconv study (**Suppl. Table 1**) were accessed through deconvData v3^6^. This new version of the deconvData collection contains an improved version of the *Morandini* dataset with refined cell-type annotations (with dendritic cells separated from monocytes), and the recently released *Dietrich* dataset^7^. Deconvolution methods were run through deconvBench v1.1.0, unless otherwise stated, while Rectangle v1.4.2 was run locally on an Apple MacBook M1 laptop using the default parameter settings.

We evaluated deconvolution performance using two metrics: Pearson’s correlation and root mean square error (RMSE). For both metrics, method estimates were compared with the available ground-truth cell-type fractions. Metrics were computed across all samples and cell types to obtain a global performance measure, and separately for each cell type to assess cell-type-specific performance. For each sample *s*, we calculated sample-wise RMSE by comparing the vector of ground-truth cell-type fractions with the corresponding vector of estimated fractions from each deconvolution method. Both vectors had length C, corresponding to the number of cell types as 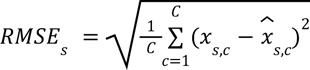. Similarly, we calculated cell-type-specific RMSE by comparing the ground truth and estimated fractions for each cell type *c* across all available samples *S* as 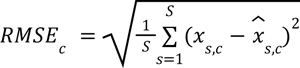

To evaluate how scRNA-seq reference size affects computational requirements, we subsampled the full *Hao* dataset (147,391 cells) into references of 1,000, 5,000, 10,000, 25,000, 50,000, 100,000 and 147,391 cells. Downsampling was stratified by cell type to preserve the cell-type proportions of the full reference. Each reference was used as input to the deconvolution methods together with the *Finotello* bulk RNA-seq dataset to be deconvolved. For each reference size, we measured runtime and peak memory use to assess how these requirements increased with the number of reference cells. This evaluation was run on an Apple MacBook M1 with 8 cores and 32 GB RAM. Among the evaluated methods other than Rectangle, only BayesPrism supports multiprocessing; therefore, BayesPrism was run using all available eight CPU cores, whereas the remaining methods were executed in their single-process configuration. SCDC (v0.0.0.9000), MuSiC (v1.0.0), BisqueRNA (v1.0.5), BayesPrism (v2.2.3), DWLS (commit b73dd13), AutoGeneS (v1.0.4), Scaden (v1.1.2), and CIBERSORTx (sha256:9dc06b0a) were run with their native implementations, using R, Python, or Docker, depending on their software requirements. For these methods, we used default settings whenever possible and applied lighter configurations only when required by runtime or memory constraints (**Suppl. Table 10**). Rectangle was run with default parameter settings. Runtime was measured as wall-clock time using Python’s `time.perf_counter()`. Peak memory use was measured with the psutil (https://pypi.org/project/psutil/) Python library by recording the maximum resident set size during each run, including child processes. For Rectangle’s scalability analysis (**Fig. 2D**), reference size, runtime, and peak memory use were expressed relative to the smallest reference size (1,000 cells) and plotted across increasing reference sizes.

To quantify Rectangle’s spillover, we leveraged the deconvBench spillover analysis^6^. Rectangle was run locally on an Apple MacBook M1 laptop with unknown content correction disabled. For the other methods, we directly considered the results obtained in the original publication^6^.

For the assessment of fine-grained deconvolution, we integrated Rectangle (run with default parameter settings) directly into the deconvBench framework to ensure the same annotations and simulated samples across all methods, leveraging the *Lambrechts* and *Wu* scRNA-seq datasets. For each dataset, five replicates of ten pseudobulk samples were generated with SimBu^3^ using the “mirror_db” scenario, which preserves cell-type proportions resembling those observed in the original scRNA-seq reference. Simulations were rerun using the fine-level cell-type annotations listed in **Suppl. Table 5**. For each method, RMSE was calculated separately for each fine cell type by comparing the estimated and true cell-type fractions across pseudobulk samples, after matching shared sample and cell-type labels. RMSE values were averaged across the five simulation replicates, and methods were then ranked within each fine cell type, with the lowest RMSE receiving rank 1, indicating the top-performing method.

### Deconvolution of spatial transcriptomics data

Transcriptomics data from the mouse brain^8^ were downloaded from ArrayExpress (https://www.ebi.ac.uk/biostudies/arrayexpress), with accession E-MTAB-11114 (10x Visium spatial transcriptomics, slide “ST8059048”) and E-MTAB-11115 (single-nuclei RNA-seq, samples “5705STDY8058280”, “5705STDY8058281”, and “5705STDY8058282”). Cell-type labels were based on the associated “annotation_1” metadata field; unknown/low-quality cells were removed, and some of the cell labels were aggregated as indicated in **Suppl. Table 11**. Spots with <500 total counts were removed. Gene counts from the spatial transcriptomic data were normalized to CPM. Rectangle was run with default parameter settings to deconvolve the cellular composition of each spot, informed by the preprocessed single-nuclei RNA-seq data. After removal of disconnected spots, Rectangle estimates were added to the Visium SpatialData^9^ object and visualized with spatialdata-plot, enabling direct overlay of the deconvolution results on the spatial coordinates. For the spatial panel, selected excitatory neuronal populations (Ext_L23, Ext_L25, Ext_L56, Ext_L6B, Ext_Hpc, Ext_Pir, Ext_Thal, and Ext_ClauPyr) were visualized by assigning each spot to the population with the highest estimated fraction and overlaying the resulting labels on the Visium spot geometry.

For systematic benchmarking of spatial deconvolution, we leveraged the “STARMap”, “seqFISH+ ob”, and “seqFISH+ cortex” validation datasets from the Spotless v1.0 benchmarking workflow^10^. We integrated Rectangle into the Spotless benchmark workflow as an additional dockerized method, while leaving the benchmark datasets, gold-standard definitions, and evaluation procedures unchanged. For Rectangle, the spatial count matrices were converted to CPM before deconvolution analysis. We ran Rectangle on the three gold-standard datasets using default parameters and the original Spotless Nextflow workflow: `nextflow run main.nf --methods rectangle -profile local,docker --mode run_standard --standard gold_standard_X -c standards/standard.config`

where ‘`gold_standard_X’` denotes the specific gold-standard datasets. Downstream performance assessment was then performed with the Spotless evaluation workflow:

`nextflow run subworkflows/evaluation/evaluate_methods.nf -profile local,docker --sp_input “standards/gold_standard_X/*.rds”`

For gold standard 3, we ran the version with 12 cell types. The Spotless repository also contains precomputed results for the previously evaluated state-of-the-art methods. These method outputs were reused as input to the same ‘`evaluate_methods.nf’` workflow and served as the reference results against which Rectangle was compared. To align with the bulk benchmarking, we considered Pearson’s correlation and RMSE as main evaluation measures.

### Analysis of The Cancer Genome Atlas RNA-seq data

For the analysis of lung cancer TCGA data, we used the *Lambrechts* single-cell RNA-seq dataset obtained via deconvData v3 as a reference dataset for signature building. We used cell-type annotations with medium resolution, with two adaptations: conventional CD4^+^ T cells and regulatory T cells (T_regs_) were aggregated into a single CD4^+^ T-cell category, and normal epithelial cells were excluded due to the difficulty in distinguishing them from malignant epithelial cells based solely on gene expression profiles. The final cell-type labels were: B cells, endothelial cells, macrophages, mast cells, monocytes, natural killer (NK) cells, neutrophils, plasma cells, stromal cells, CD4^+^ T cells, CD8^+^ T cells, tumor cells, myeloid dendritic cells (mDCs), and plasmacytoid dendritic cells (pDCs). Ribosomal genes were removed from the single-cell reference. Based on this data, we generated two references for signature building, either including or excluding tumor cells, and used them to run Rectangle with default parameters on TCGA lung cancer samples (n=1,012), thereby evaluating deconvolution when tumor signal was explicitly represented or captured as unknown content. Consensus tumor purity estimates (“CPE”) for the TCGA samples were obtained from a previous study^11^.

For the deconvolution of breast cancer TCGA data, Rectangle was trained using the *Wu* single-cell dataset, accessed from deconvData v3. We used the “celltype_minor” annotation level and excluded cycling cells (Cycling T-cells, Cycling Myeloid, Cancer Cycling, Cycling PVL) and normal epithelial subpopulations (Myoepithelial, Luminal Progenitors, Mature Luminal). Rectangle was run with default parameter settings to deconvolve TCGA bulk RNA-seq samples (n=1,091). For downstream analysis, Rectangle deconvolution results were combined with cancer-immune phenotype annotations for TCGA breast cancer samples^12^, obtaining matched data across 1,068 patients: (D, n=400), fibrotic (F, n=215), immune-enriched fibrotic (IE/F, n=180), and immune-enriched (IE, n=273). Aggregated cell-type fractions were derived by summing the corresponding minor cell-type estimates as indicated in (**Suppl. Table 12**). Pairwise differences between immune-subtype groups were assessed using Wilcoxon rank-sum test, with Benjamini-Hochberg correction for multiple testing applied globally across all plotted cell types and pairwise subtype comparisons.

### Analysis of developing brain organoids

The scRNA-seq and bulk RNA-seq data from human brain organoids^13^ were retrieved from the Gene Expression Omnibus (GEO, https://www.ncbi.nlm.nih.gov/geo/), with accession numbers GSE277968 and GSE277968, respectively. Samples from time-series organoid data were selected, gene counts in the bulk dataset were assigned to gene names (from Ensembl identifiers), and normalized to CPM. For signature building and deconvolution, the coarse annotation of the scRNA-seq data was considered. Rectangle was run with default parameter settings. For downstream evaluation, sample metadata were parsed to extract replicate, cell line, protocol, and differentiation day. Dorsal-protocol H9 and 176 samples were compared over time for selected cell types and unknown content. Unknown content was quantified using the ‘Unknown’ fraction estimated by Rectangle for each bulk sample. Samples were grouped by protocol, cell line, and differentiation day, and the unknown fraction was summarized across biological replicates to assess how the unexplained bulk signal changed over organoid development. To investigate transcriptional programs associated with this unknown component, we used Rectangle’s gene-level reconstruction error *e _g_*, which measures the discrepancy between the observed and reconstructed bulk expression profile for each gene and sample: *e_g_ = y_g_ − ŷ_g_*, where *y _g_* is the observed bulk expression of gene *g*, and ŷ*_g_* is the reconstructed expression of that gene from the signature matrix and estimated cell fractions. At the earliest time point, reconstruction errors were averaged across replicates within each protocol and cell-line group. For each group, the 10 genes with the highest mean reconstruction error were selected, and genes were then ranked by how frequently they appeared across these top-10 lists. This recurrence-based summary was used to identify genes and transcriptional programs that consistently contributed to the unexplained component across different protocols and cell lines.

### Overall assessment and visualization

To provide an integrated overview of method performance, we summarized all benchmark results in a single heatmap-like graph, in which each row corresponds to a deconvolution method and each column to a performance metric evaluated on a given dataset. The columns were drawn from five evaluation categories. *Accuracy* contributed two metrics per dataset: the root mean square error (RMSE) and the correlation between estimated and ground truth cell type proportions, across the six gold-standard bulk RNA-seq datasets. For each dataset, both statistics were computed cumulatively across all individual cell types. *Efficiency* contributed two columns: the runtime and the peak memory usage measured on the subsampled 20k-cell *Hao* dataset. *Spillover* was represented by three columns, one for each spillover dataset, reporting the spillover percentage. *Resolution* contributed the cumulative RMSE across all cell types for each of the two fine-grained pseudobulk datasets (*Wu* and *Lambrechts*). Finally, robustness to unknown content contributed two metrics: the “all cells” RMSE and correlation values computed across all pseudobulks considered for **Fig. 5A-B**. Replicate-level values were averaged within each dataset and unknown-content fraction, and the heatmap used the mean of these all-cells summary values per method.

To make metrics comparable across their different scales and directions, each column was independently rescaled in [0, 1] by min-max normalization, excluding missing values. For metrics in which lower values indicate better performance (RMSE, runtime, memory, and spillover), the scale was inverted, preserving the convention that 1 denotes the best and 0 the worst method for every metric. To rank methods, we combined the normalized metric scores into an overall score per category. We first computed a per-category score for each method as the mean of its min-max normalized scores within that category; missing values were considered as 0. The final score was computed as the sum of the five category scores and used to rank methods.

