## Supplementary Figures for "Rectangle: robust and scalable multiscale deconvolution informed by single-cell RNA sequencing data"

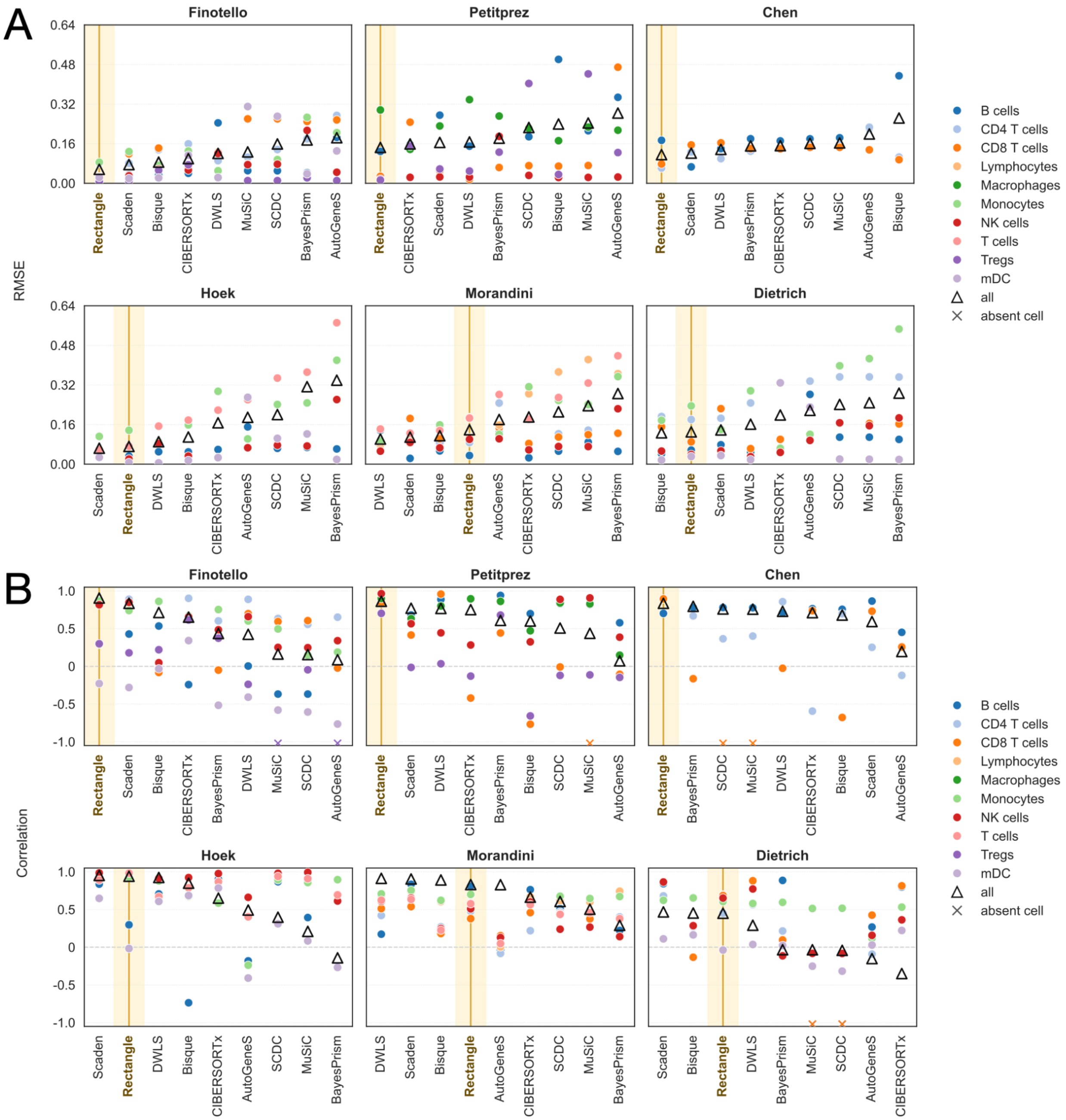


**Suppl. Fig. 1 | (A)** Per-cell-type root mean square error (RMSE) between true and estimated cell fractions for different deconvolution methods across six benchmark datasets. Filled circles denote individual cell types (color), open triangles denote the correlation across all cell types together ('all'), and crosses identify cell types identified as absent in all bulk samples. Methods are sorted by the 'all' cell-type RMSE. **(B)** Per-cell-type Pearson correlation between true and estimated cell fractions for different deconvolution methods across six benchmark datasets. Filled circles denote individual cell types (color), open triangles denote the correlation across all cell types together ('all'), and crosses identify cell types identified as absent in all bulk samples. Methods are sorted by the 'all' cell-type correlation..

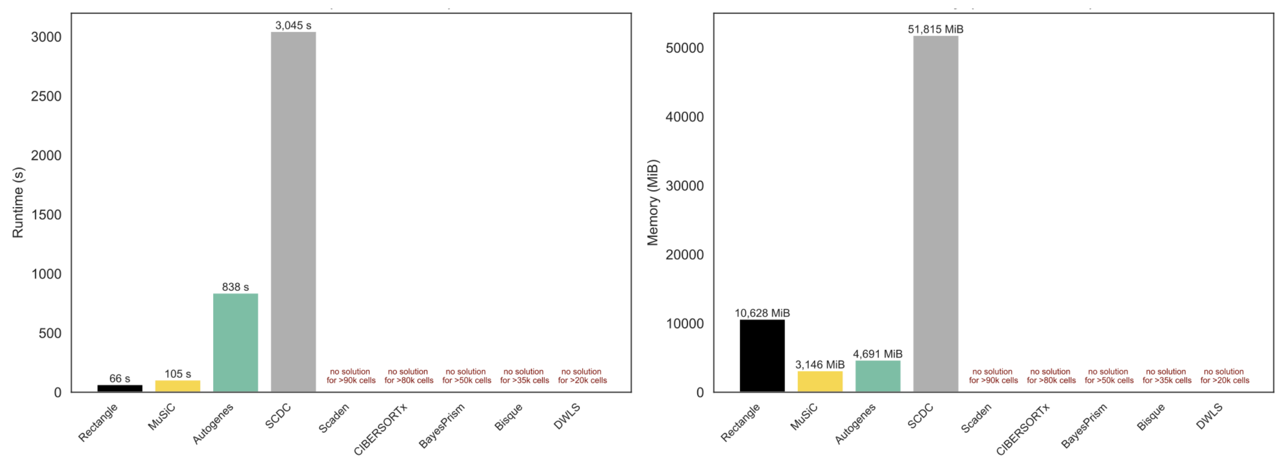


**Suppl. Fig. 2 |** Runtime in seconds **(**left**)** and peak memory usage in MiB **(**right**)** for each method on the full *Hao* reference. Methods are sorted left-to-right by increasing runtime. Methods that could not be run on the full reference are indicated with red text.

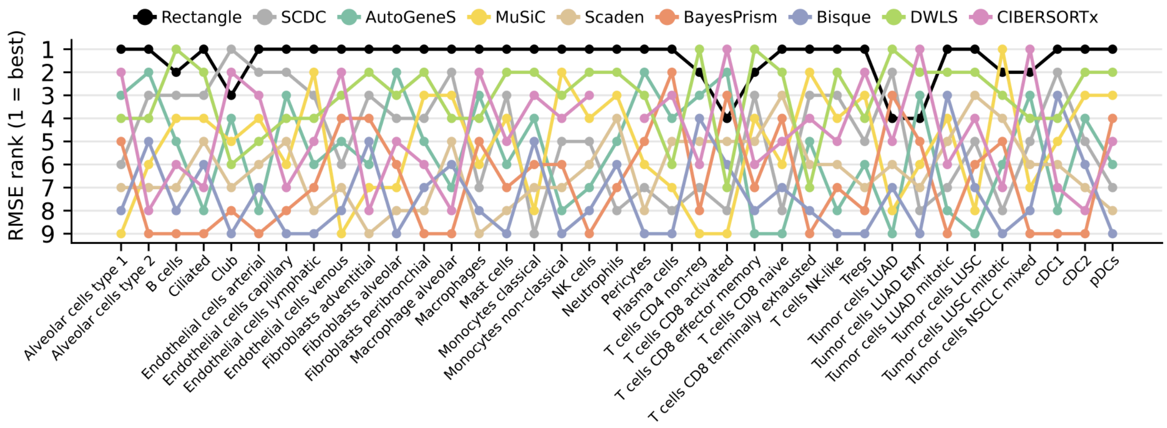


**Suppl. Fig. 3 |** Cell-type ranks based on root mean square error (RMSE) of each method across 49 fine-grained cell types from the *Lambrechts* dataset. The top-performing method for each cell type appears at the top of the bump chart (rank=1).

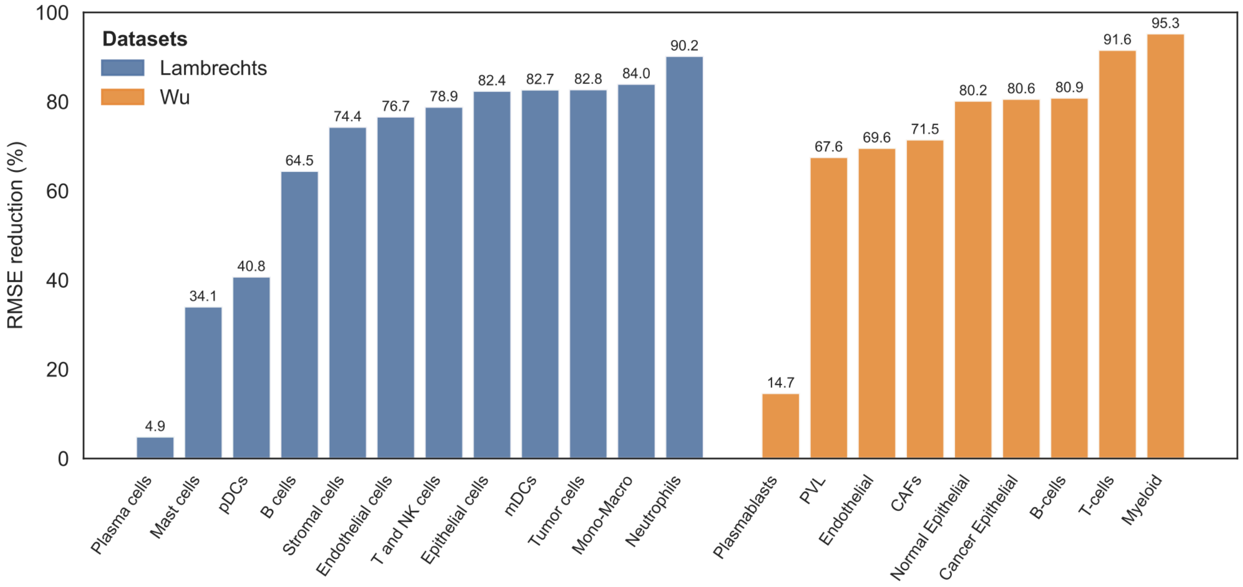


**Suppl. Fig. 4 |** Bar chart showing the per-cell-type RMSE reduction (%) when deconvolving with fine-grained annotations and aggregating to coarse cell types, compared to direct coarse-level deconvolution. Results are shown for the Lambrechts (lung cancer, blue) and Wu (breast cancer, orange) datasets.
